# Control and simulation impact on nonlinear Hepatitis-B model by using Padé-approximation based Differential Evolution

**DOI:** 10.1101/831636

**Authors:** Muhammad Farman, Muhammad Farhan Tabassum, Muhammad Saeed, Nazir Ahmad Chaudhry

## Abstract

Hepatitis B is the main public health problem of the whole world. In epidemiology, mathematical models perform a key role in understanding the dynamics of infectious diseases. This paper proposes Padé approximation (*Pa*) with Differential Evolution (*DE*) for obtaining solution of Hepatitis-B model which is nonlinear numerically. The proposed strategy transforms the nonlinear model into optimization problem by using Padé approximation. Initial conditions are converted into problem constraints and constraint problem become unconstraint by using penalty function. *DE* is obtained numerical solution of Hepatitis-B model by solving the established problem of optimization. There is no need to choose step lengths in proposed Padé-approximation based Differential Evolution (*PaDE*) technique and also *PaDE* converges to true steady state points. Finally, a convergence and error analysis evidence that the convergence speed of *PaDE* is greater than Non-Standard Finite Difference (*NSFD*) method for different time steps.

## Introduction

The era of statistics, biotechnology and medicine is known for the revolutions and a great amount of information which has been produced with the acceleration in the method of know-how discovery of biological structures. Such advancements are modifying the mode of biomedical studies, programs and measures of improvement are performed. Clinical information supplements organic records, allowing certain descriptions of each wholesome and diseased state, in addition to disorder progression and reaction to remedies [1]. The accessibility of records demonstrating innumerable organic conditions, tactics and their dependencies on time permits to have a look at organic structures at diverse tiers of employer from molecules to organism or even up to the population degree [2].

Human interest includes modeling which is the representation, manipulation and communication of regular real world objects of day to day life. As you possibly can effortlessly comprehend, there are numerous ways to take a look at an item or, equivalently, there are many unique observers for the same object. Any observer has ‘one of a kind perspectives’ of the identical item, i.e. ‘there’s no omniscient observer with unique get right of entry to the reality’. Each distinct observer collects facts and generates hypothesis that are constant with the records. It is a logical procedure is known as ‘abduction’. Which isn’t always dependable, although with the admiration to a systematic yet unidentified to which we are all blind [3].

The model of an outline is a method in terms of constituents’ items along with the connections amongst these items, wherein the explanation and description itself is generally easy to decode or interpret by using people [4]. At some point of the records exquisite epidemic illnesses have unfold the world over, taking human populations towards disastrous effects in future. Millions of survivors were sheltered. Cholera and Black dying are such epidemics which have been the causes of severe migration [5]. Expertise approximately the un-fold and the sever situation of epidemic infections is valued in stopping the stark damages to human population [6].

Hepatitis is known an illness, characterized as irritation or swelling on liver. Though it has several causes as constant use of alcohol and pills, yet viruses are usually the prominent causes of such epidemics. Hepatitis B is a grave threat to health mission [7]. Approximately one billion of population around the sphere was inflamed with hepatitis B virus known as HBV and over 3 hundred million people were prevented from the virus [8]. Severe instances can result in liver failure and demise. However maximum sufferers with severe hepatitis sooner or later were recovering completely. In a few patients, the ailment turns into chronic and is referred to as chronic hepatitis [9]. Human beings with persistent hepatitis may also revel in mild, indistinct symptoms of fatigue and poor urge for food. Continual hepatitis can result in a liver disease referred to as cirrhosis, and it is additionally an important motive of liver cancer [10].

Hepatitis kinds A, C, D, and E are because of viruses that have a middle of ribonucleic acid (*RNA*). Hepatitis B virus (*HBV*) is a deoxyribonucleic acid (*DNA*) core. Hepatitis B is a critical public fitness hassle that impacts human beings of all ages around the sector. An infectious virus attacks the liver and becomes the cause disease. *HBV* infection leads towards serious health issues that harms the liver and consequently causes death [11].

There are many mathematical models which are being established to model and study the dimensions and dynamics of this severe and grim epidemiologic infection [12, 13]. These models incorporated the contact amid *HBV* while dealing with both the fit and diseased cells of liver equally. Few models have studied the resistant responses which are also adaptive in combating viruses which are free and reduce the disease-ridden cells [14-17]. Adaptive immunity has been represented by antibody immune and cytotoxic T-lymphocytes (*CTL*) responses. The viral infection of *HBV* with *HBV DNA*- having capsids were categorized by mathematical analysis [18-22]. There arises a significant point that diseased cells of liver discharge *HBV DNA*-which contains capsids in the form of mature viruses after being enclosed by viral envelope proteins cellular membrane and lipids [23, 24]. Recently, the optimal control of *HBV* infection with *HBV DNA*-having capsids and *CTL* immune response had been studied [22].

These epidemiological models are vital procedures to investigate and acquire improved information about the development with the help of Mathematical tool which are built on arithmetical and numerical analysis, influence and the deriving mechanisms, particularly when there is not available any analytical solution. A thoughtful information and knowledge about these model aid in adopting preventive actions and to evaluate their efficiency and effectiveness to avert such infections.

Contemporary meta-heuristics are proposed to cope up the maximum hitches by changing them into optimization problems in recent times. As we know Meta-heuristic algorithms have been formulated by natural phenomena as swarm behaviors [25, 26], evolution [27, 28], sport strategies [29], water dynamics [30, 31], food foraging behavior [32] etc. Consult the survey article for more detailed studies referred in [33]. Meta-heuristics is based on the approaches which are resolving differential equations related to the class of non-standard mesh free methods. The improvisation of these recommended heuristics to differential equations may also found in [34, 35], but here is a problem that the applications of these Meta-heuristics to widespread and epidemic models are really hard to see.

This study presents a novel and innovative scheme known as Padé-approximation [36] which is based on Differential Evolution Algorithm to handle the numerical treatment of this model. This recommended computational framework encompasses these following features:

i. By constructing an equivalent optimization problem by manipulating the extrapolation and interpolation métiers of Padé approximation.
ii. To preserve the positivity by initial bounded-ness and conditions of agreement by outlining the problem constraints.
iii. An indispensable prerequisite by using the penalty functions approach to construct fitness/objective function
iv. The implementation of differential evolution (DE) to optimize the constructed fitness function.

This advanced method is acclaimed as Padé approximation with Differential Evolution (*PaDE*). This whole paper has been established on these grounds. Section 2 based on the fundamentals of Padé approximation and Differential Evolution. Section 3 has the comprehensive detail of Hepatitis B model. Section 4 delineates the offered structure of *PaDE* scheme to solve the numerical treatment of nonlinear Hepatitis B model. While in Sect. 5, revolves around the analyses of the results which have been presented. Finally, few concluding remarks and findings for future directions have been given.

## Numerical Results

For numerical illustrations set parameters of *DE* algorithm: *N* = 50; *F* = 0.55; *CR* = 0.91 and *maximum iterations* = 2000. Padé approximation order is set as (*N, M*) = (2, 2). The parameter q_max_ is set as 2000. Penalty factor is set to be Lq = 10^10^ for all q. The optimized parameters of Hepatitis B model are given in Table 1. The mathematical analysis of epidemic Hepatitis B model with non-linear occurrence has been offered with parameter values in table 1. To notice the sound effects of the *PaDE* algorithm on susceptible, infected and recovered population comparison with *NSFD* having the property of uniqueness and positivity represent in figure 1-3. Figs. 4-9 show convergence solution with relationship between the different population compartments for diseases free equilibrium by using *PaDE* algorithm and NSFD scheme, here it can be easily observe that the results *PaDE* algorithm are more reliable and better convergence as with numerical scheme. In figure 8 and 9 represents the impact of vaccination on infected and chronic population which decreases rapidly. It also shows the susceptible and recovered population increase in time *t* after measuring the impact of vaccinations the disease should be easily controlled.

**Table 1.**
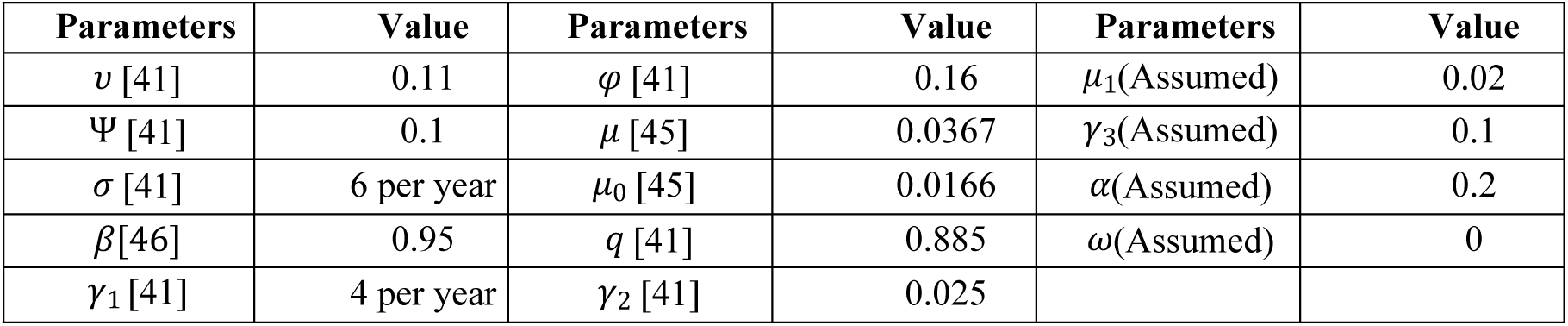
Values of physical parameters of the model.

**Figure 1.** Dynamical behavior for susceptible population S (t) in a time *t* with PaDE and NSFD.

**Figure 2.** Dynamical behavior for infected population I(t) in a time *t* with PaDE and NSFD.

**Figure 3.** Dynamical behavior for recovered population R(t) in a time *t* with PaDE and NSFD.

**Figure 4.** Dynamical behavior for S(t),C(t),R(t) populations in a time *t* with PaDE and NSFD.

**Figure 5.** Dynamical behavior for S(t),C(t),V(t) populations in a time *t* with PaDE and NSFD.

**Figure 6.** Dynamical behavior for S(t),I(t),R(t) populations in a time *t* with PaDE and NSFD.

**Figure 7.** Dynamical behavior for L(t),I(t),C(t) populations in a time *t* with PaDE and NSFD.

**Figure 8.** Dynamical behavior for S(t),V(t),C(t),R(t) in a time *t* with PaDE and NSFD.

**Figure 9.** Dynamical behavior for S(t),V(t),I(t),R(t) in a time *t* with PaDE and NSFD.

## Error Analysis and Convergence Analysis with PaDE

In Fig. 10-15 represents error analysis of different population compartments of the developed algorithm in a time *t* with step size *h* = 1 and *h* = 2 and for disease free equilibrium points. It can be easily seen that by reducing the step size the model (1) converge rapidly to the steady state point.

**Figure 10.** Error Analysis of S(t),C(t),R(t) to PaDE Algorithm at step size h=1,h=2.

**Figure 11.** Error Analysis of S(t),I(t),R(t) to PaDE Algorithm at step size h=1,h=2.

**Figure 12.** Error Analysis of S(t),C(t),V(t) to PaDE Algorithm at step size h=1,h=2.

**Figure 13.** Error Analysis of L(t),I(t),C(t) to PaDE Algorithm at step size h=1,h=2.

**Figure 14.** Error Analysis of S(t),V(t),I(t),R(t) to PaDE Algorithm at step size h=1,h=2.

**Figure 15.** Error Analysis of S(t),V(t),C(t),R(t) to PaDE Algorithm at step size h=1,h=2.

## Discussion

A mathematical model for Hepatitis B transmission has been shown with variable aggregate population size and distinctive transmission rate in populations. It has been verified existence of an ailment free equilibrium and processed the fundamental reproduction number *R*_0_ utilizing the technique, where it is asymptotically stable and sensitivity analysis of the parameters include in threshold parameter *R*_0_. This study proposed Padé approximation based differential evolution (*PaDE*) algorithm for solution of a nonlinear Hepatitis B model. For treatment of Hepatitis B model new alliance of Padé approximation and *DE* was developed. Great accuracy of developed algorithm satisfies the governing equations. Through constraints and the penalty function all the conditions of the solution were efficiently handled. The acquired solution retains very rapid convergence and superior to *NSFD*. The comparison of the above figures validates that the solutions from *PaDE* algorithm is in good agreement with *NSFD* particular. By using its higher order and more robust optimization approach the accuracy of the numerical solution can be enhanced. It has been demonstrated systematically and explained by numerical reproductions that our *PaDE* algorithm is vigorously steady and comparison with *NSFD* scheme regarding the accompanying properties of the continuous model: positivity and boundednes of solution, local and global stability of equilibrium. It is essential to take note of that *PaDE* algorithm for mathematical models dependent on system of differential equation is all the more incredible way to deal with process the convergent solutions and error analysis for the ailment models. We introduced the numerical simulation and confirmed all the logical outcomes numerically by developed algorithm to diminish the contaminated rates quick for sickness free equilibrium by utilizing distinct step sizes.

## Material and Methods

### Padé approximation

Through classical theory of continued fractions the idea of a Padé-approximation was introduced at the end of the 19th century. The (*N, M*) order rational function of Padé approximation is [37]

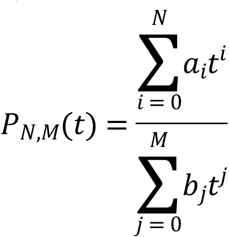

The polynomials 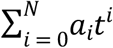 and 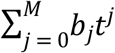 are known as Padé approximants. By putting *b*_0_ ≠ 0 normalizing the above expression and attain the following form:

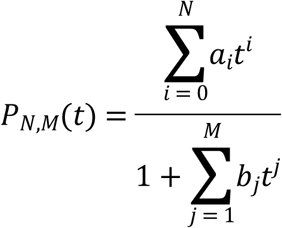

The above expression contains (*N* + *M* + 1) undetermined coefficients, applying the Maclaurin series expansions of *P*_*N,M*_(*t*) to get the target referred in [36].

### Differential evolution

In 1995 Storn and Price introduced stochastic population based algorithm named as Differential Evolution (*DE*) [38], it was simple, robust and reliable to find optimal point. The *DE* strategy [39] which the most often used is implemented in this paper. The individual set of parameters represented by *D* dimensional vector. *NP* represent member in population with vectors 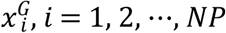 and *G* indicates a generation. During the evolution process the generation remains same. Randomly choose the initial population with uniform distribution. After initialization *DE* has following three operators:

#### 3.1. Initialization

Initial population must be generated for optimization procedure and assigned a randomly chosen value from the boundary constraints:

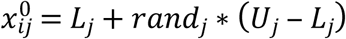

where *rand*_*j*_ indicates a uniform distribution between [0, 1], *L*_*j*_ is lower and *U*_*j*_ is upper bounds for the j^th^ decision parameter [28].

#### 3.2. Mutation

A mutant vector namely *v* is produced for each target vector 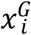:

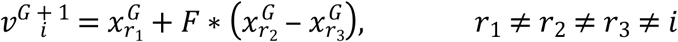

with randomly chosen indices and *r*_1_,*r*_2_,*r*_3_ ∈ {1, 2, …, NP}.

*F* ∈ ℝ which control the amplification of 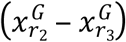, the value of F must be within the range of [0, 2] [28]. The above indices must be different from each other and also different from the running index *i* so *NP* at-least four. The new value of this component is generated using if component of a mutation vector exceeds the bounds of search space.

#### 3.3. Crossover

To yield the trial vector namely *u* the following scheme is used, the target vector is mixed with the mutated vector,

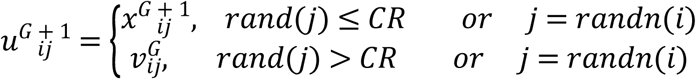

where *j* = 1, 2, …, *D, rand*(*j*) ∈ [0, 1] is the j^th^ evaluation of a generator number. Crossover probability constant is *CR* ∈ [0, 1]. *randn*(*i*) ∈ {1, 2, …, *D*} is an index which has been randomly chosen, that index confirms that 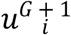 develops at least one element from 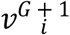.

#### 3.4. Selection

The selection scheme is as follows:

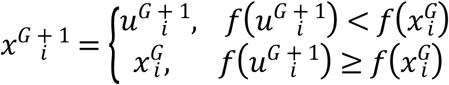

If trial vector 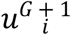 has a better fitness function value than 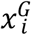, then 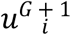 is set to 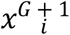. Otherwise, the old value 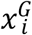 is retained due to greedy selection strategy of *DE*.

#### Constraints Handling

To handle constraints the most effective method has been used in the form of penalty function referred in [40]. In penalty function a large positive number depending on degree of violation of constraints is added to the objective function. In the following relation, objective function presented by *ψ*(***x***) and the penalty function is presented by *ζ*(*x*) describes penalized function *φ* (*x*) which was unconstrained defined as follows:

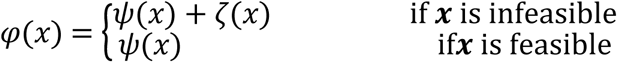

Here *ζ*(*x*) ≥ 0 is used for minimization problem and *ζ*(*x*) ≤ 0 for a maximization problem. There is very rare existence of unconstrained real world problems generally DE is designed for unconstrained optimization problems. So this technique converting the constrained optimization problems into unconstrained optimization problems.

## Mathematical model of Hepatitis B Virus (HBV)

The variables of the model at any time *t* are defined as:

*S*(*t*): susceptible individuals; *L*(*t*): latent individuals; *I*(*t*): infected individuals; *C*(*t*): chronic carriers individuals; *V*(*t*): vaccinated individuals and *R*(*t*): recovered individuals.

The model parameters are:

*μ* = birth rate; *ω* = proportion of births without vaccination; *υ* = proportion of births vertically infected; Ψ = rate of waning vaccine-induced immunity; *μ*_0_ = natural mortality rate; *β* = transmission coefficient; *φ* = reduced transmission rate relative to acute infection by carriers; *γ*_3_ = vaccination rate of the susceptible individuals; *σ* = rate of moving from latent state to acute state; *γ*_1_ = rate of moving from acute to other compartments; *q* = average probability that an individual fails to clear an acute infection and develops to carrier state; *qγ*_1_ = rate of moving from acute to carrier; *μ*_1_ = HBV-related mortality rate; *γ*_2_ = rate of moving from carrier to immune; 1 − *ω* = proportion of births vaccinated; (1 − *q*)*γ*_1_ = rate of moving from acute to recovered class.

The model under consideration was purposed by [41] under the form of the following nonlinear system of differential equations:

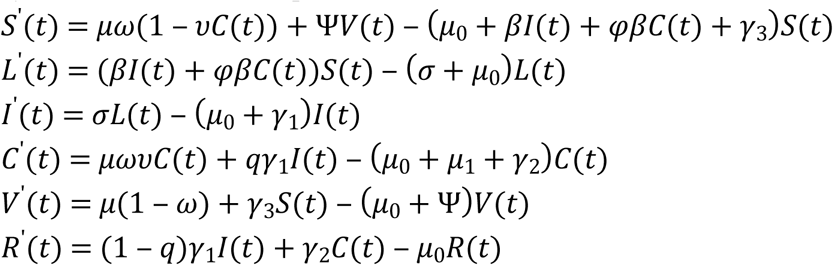

The model has been modified by taking chronic carriers are treated at the rate α [42], the newborns to carrier mothers infected at birth [41] and treated individuals recover [43, 44]. The model with system of governing equations is given as:

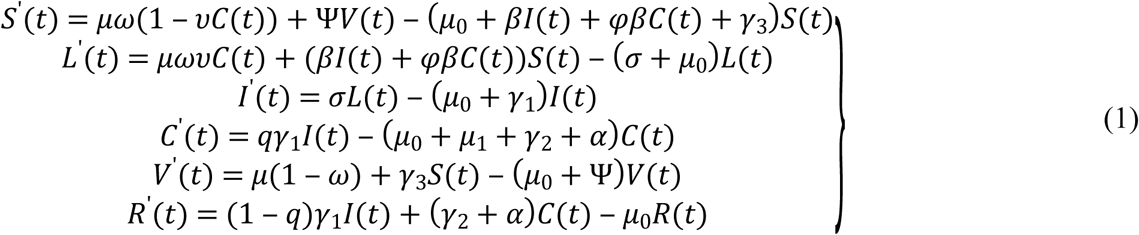

Here

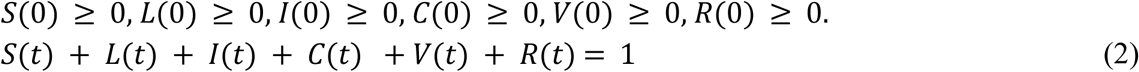

Subject to the conditions

*S*_0_ = *S*(0) = 0.7, *L*_0_ = *L*(0) = 0.05, *I*_0_ = *I*(0) = 0.05, *C*_0_ = *C*(0) = 0.08, *V*_0_ = *V*(0) = 0.06, *R*_0_ = *R*(0) = 0.12

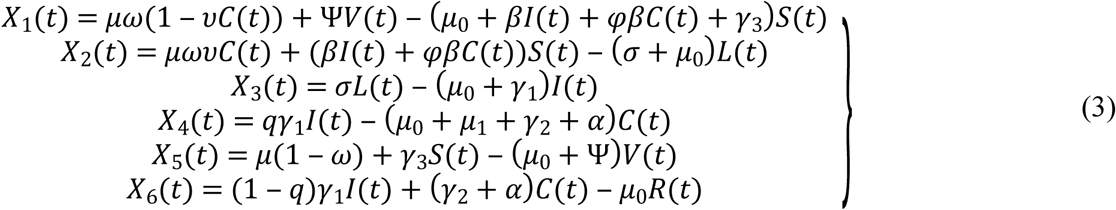

To evaluate the equilibrium point, we take

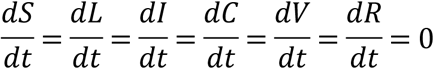

Above model becomes

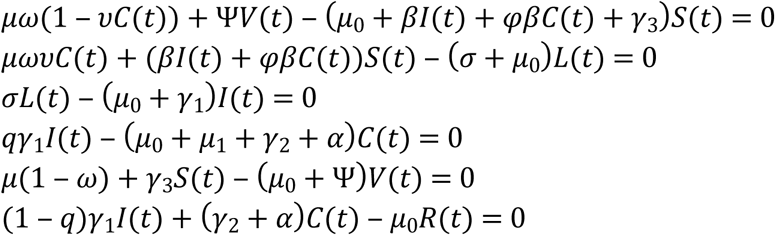

By using parametric values from Table 1, we get

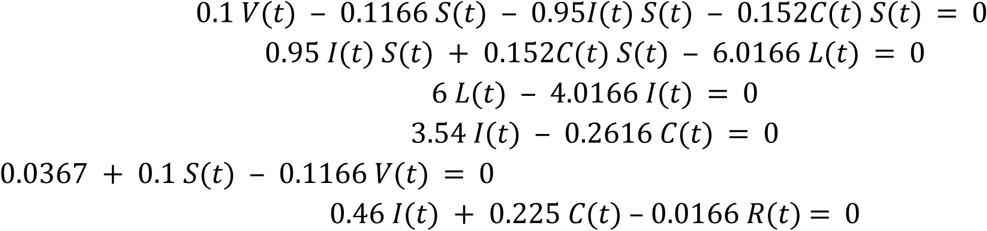

By solving all the above equations

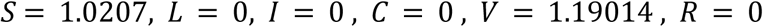

The average number of secondary infections produced by a primary infection can be identified through reproductive number. The basic reproduction wide variety R_0_ of the version is referred in [53]

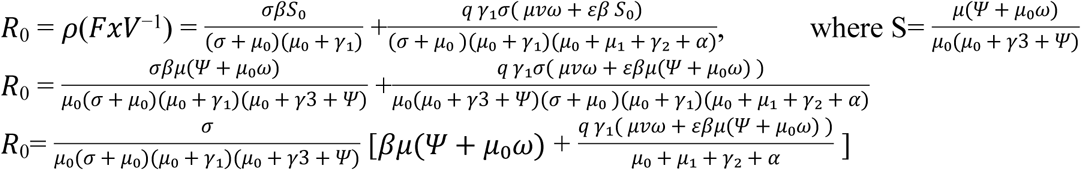

Disease Free point is

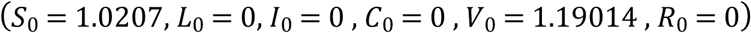

After simplification in form of parameter, the endemic point, *R*_0_ = 0.76200 < 1

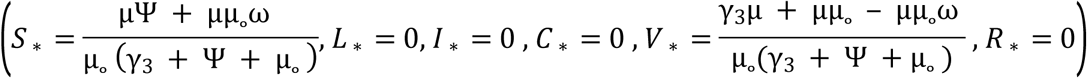

### Stability Analysis

*E*_0_ is locally stable if *R*_*e*_(*λ*) < 0, otherwise unstable.

Consider a Jacobian matrix

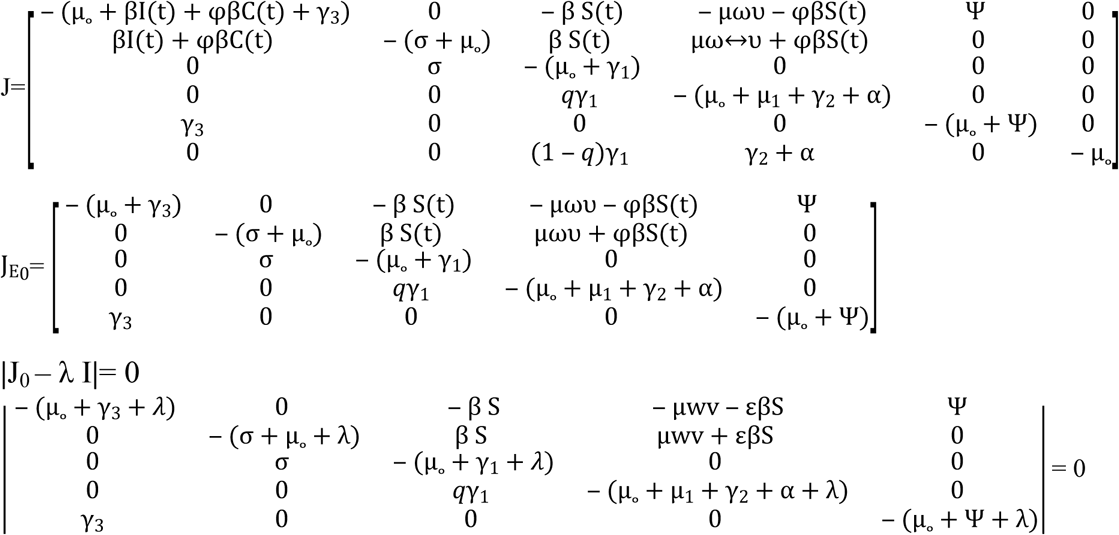

After implication and putting parameter values, the Eigen values that is;

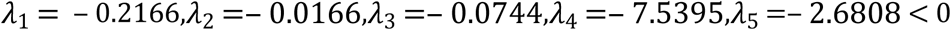

All the Eigen values are negative which shows that the given system is locally stable.

## Padé-approximation based Differential Evolution Strategy

The design of the proposed Padé-approximation based Differential Evolution involves the following main steps.

### Padé approximation based residual function

Suppose that *S*(*t*), *L*(*t*), *I*(*t*), *C*(*t*), *V*(*t*) *and R*(*t*) are approximated by Padé rational functions as

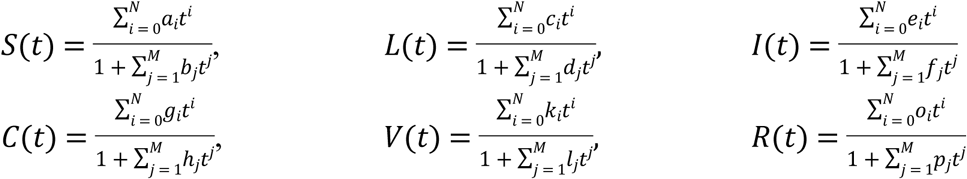

Imposing initial conditions *S*(*t*_0_) = *S*_0_, *L*(*t*_0_) = *L*_0_, *I*(*t*_0_) = *I*_0_, *C*(*t*_0_) = *C*_0_, *V*(*t*_0_) = *V*_0_ *and R*(*t*_0_) = *R* we obtain

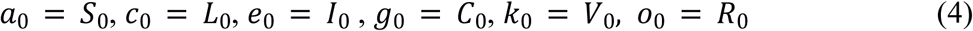

The discrete time steps are *t*_*q*_ = *t*_0_ + *qh; q* = 0, 1, 2, 3, …, *q*_*max*_, and the above system of equations (3) reduces as:

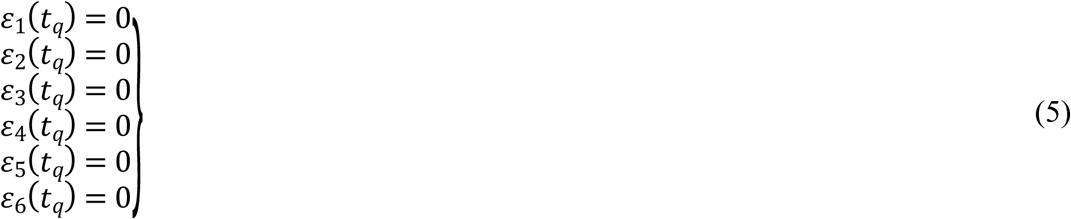

Here *ε*_1_, *ε*_2_, *ε*_3_, *ε*_4_, *ε*_5_ and *ε*_6_ are the residuals defined by

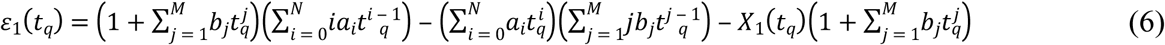

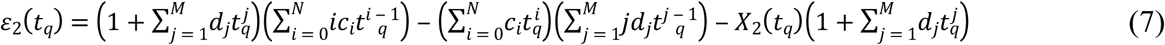

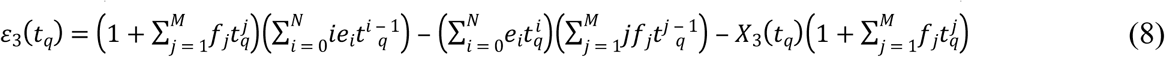

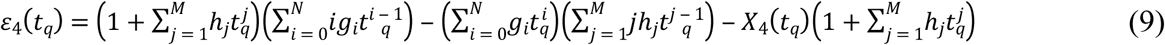

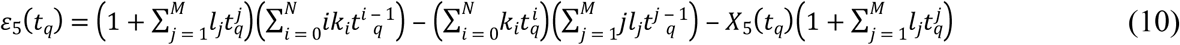

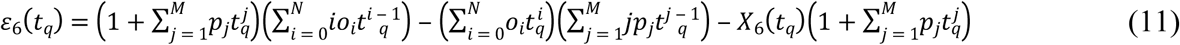

By solving system (5) having 6*q*_*max*_ nonlinear simultaneous equations the problem reduces to finding 6(M + N) coefficients of Padé approximants.

### Objective Function

Suppose that

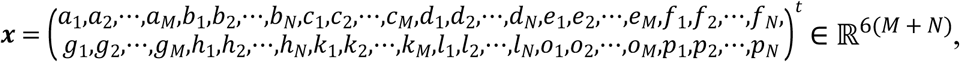

by converting the system (5) into minimization problem as:

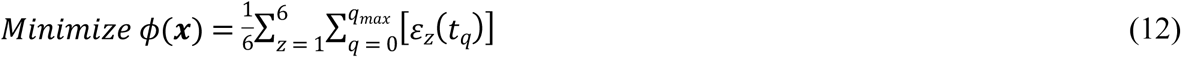

### Problem Constraints

The equality constraints of the model are considered as stated in system (4):

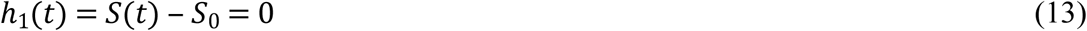

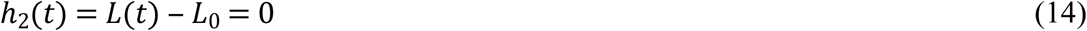

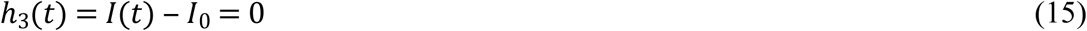

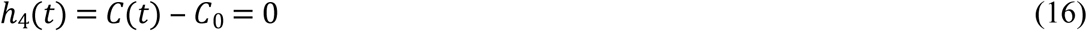

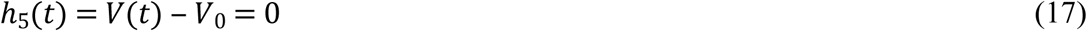

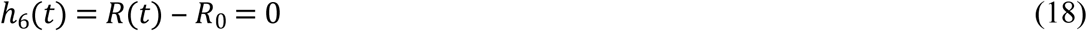

The inequality constraints (19) to (24) are related to positivity

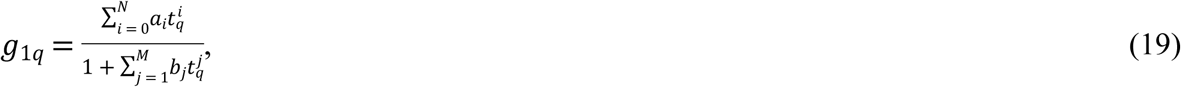

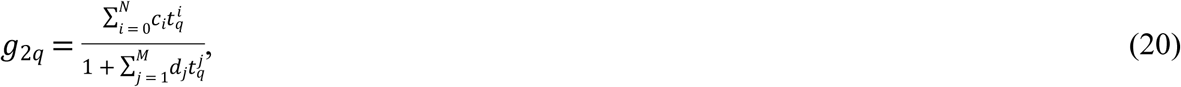

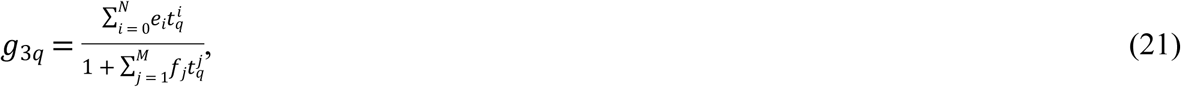

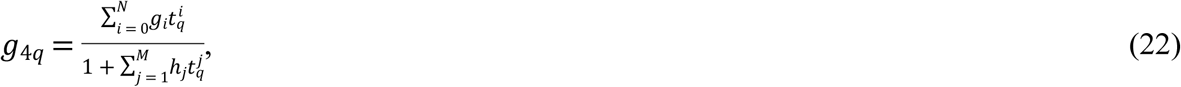

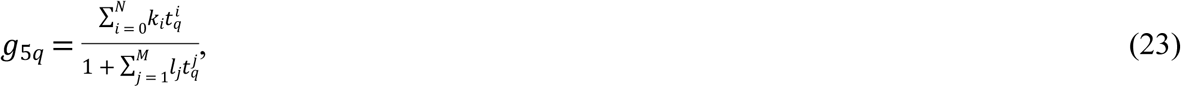

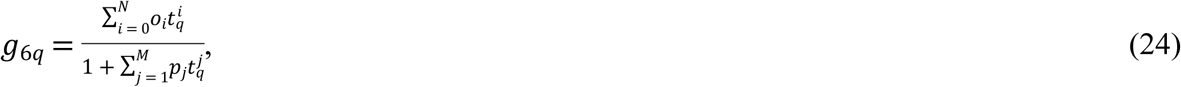

whereas (25) incorporates the bounded-ness of the numerical solution.

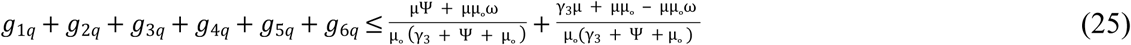

### Penalty Function

By using the following penalty function methodology unconstrained optimization model is obtained:

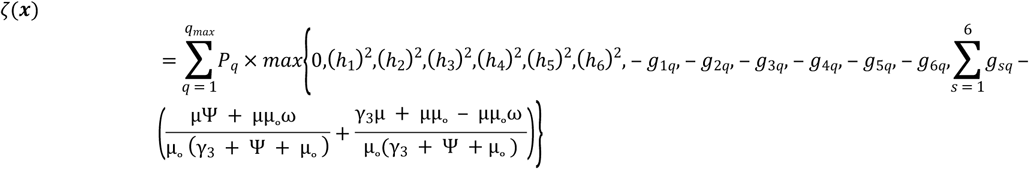

Here scalar *Pq* is a large positive real number of q^th^ discrete time step acting as a penalty factor then the unconstrained objective function is

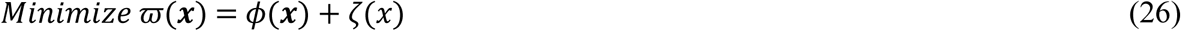

### Optimization process with differential evolution

The following steps are involved to optimize objective function (26) through *PaDE* scheme as:

Step 1. Generate population randomly, population of *K* solutions *x*_*j*_ ∈ ℝ^6(*M* + *N*)^;1 ≤ *j* ≤ K.

Step 2. Evaluate the value 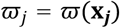 of each solution. Collect the best solution with the minimum value of objective function. Initially set T = 0.

Step 3. Set T = T + 1.

Step 4. Choose three distinct solutions ***x***_*A*_, ***x***_*B*_ and ***x***_*C*_ from the population excluding ***x***_*j*_ for each of *j* = 1, 2, 3, …, *K*, Set **y = *x***_***j***_.

Step 5. For each of the dimensions *i* = 1, 2, 3, …, 6(*M* + *N*), alter the i^th^ coordinate according to

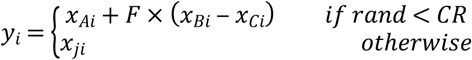

Step 6. If 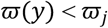 then *x*_*j*_←*y*, otherwise discard *y*.

Step 7. Best solution must be update.

Step 8. If T > number of iterations, then terminate, by maintaining the best solution, otherwise repeat all the process from step 3.

**Figure 16.**
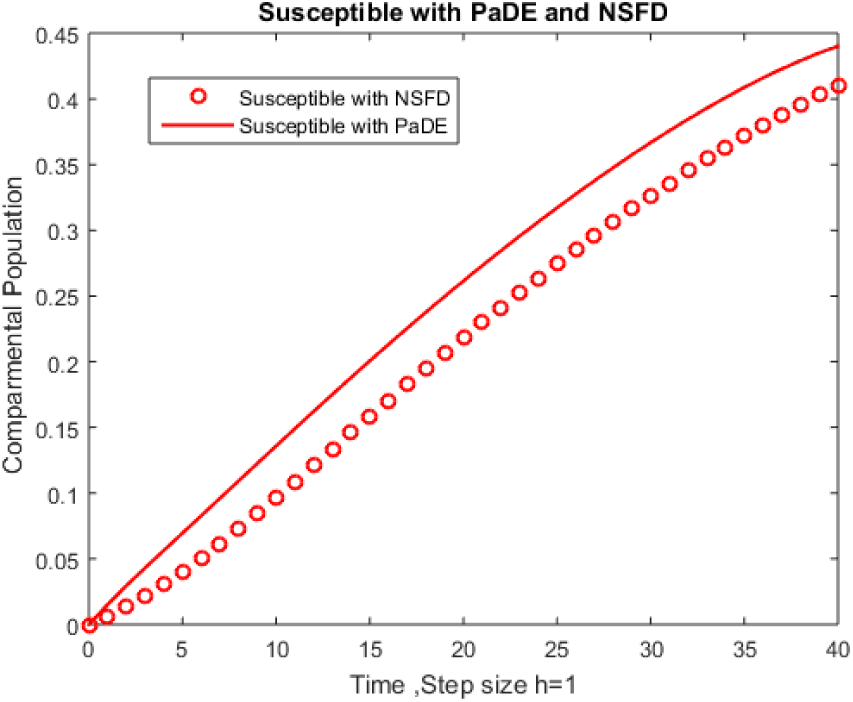
Dynamical behavior for susceptible population S (t) in a time *t* with PaDE and NSFD.

**Figure 17.**
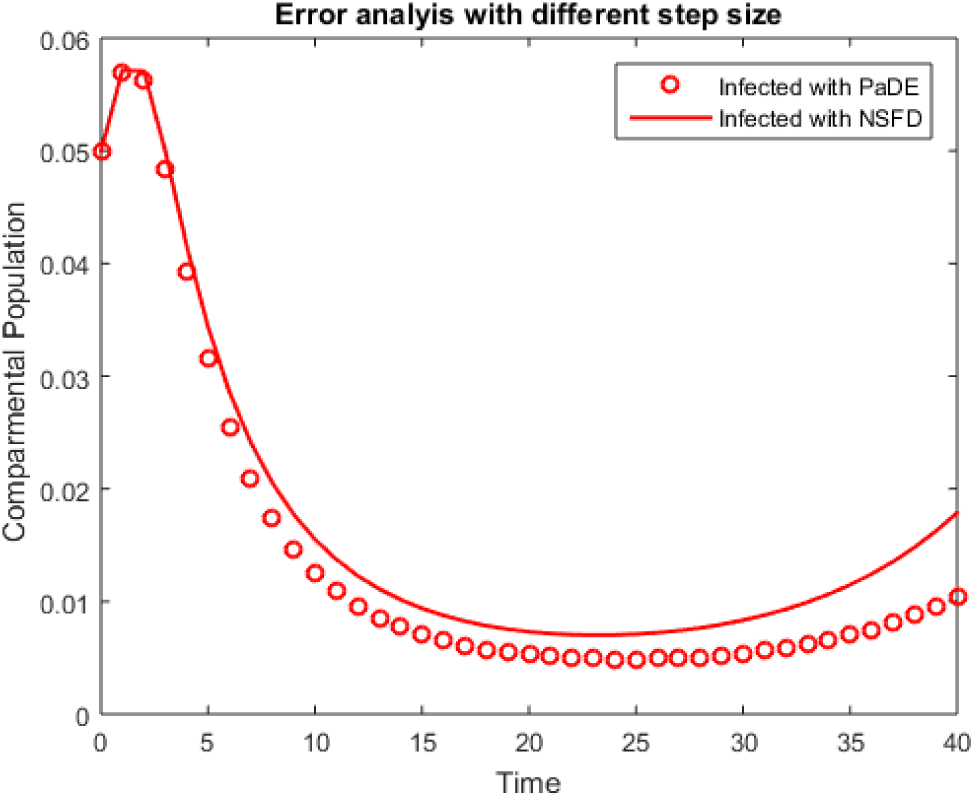
Dynamical behavior for infected population I(t) in a time *t* with PaDE and NSFD.

**Figure 18.**
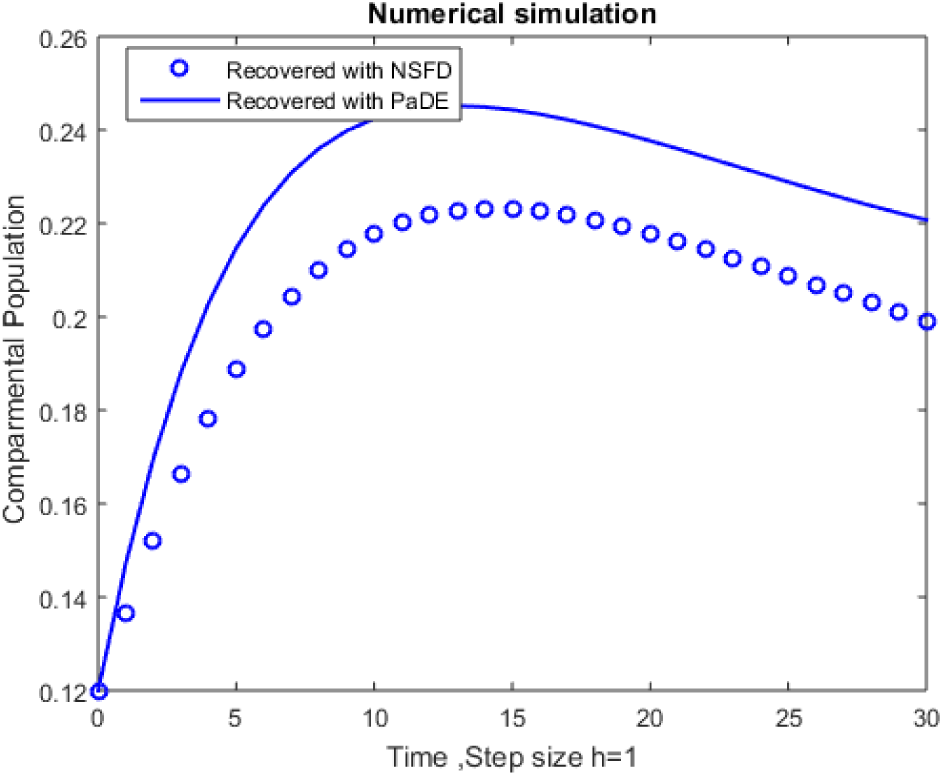
Dynamical behavior for recovered population R(t) in a time *t* with PaDE and NSFD.

**Figure 19.**
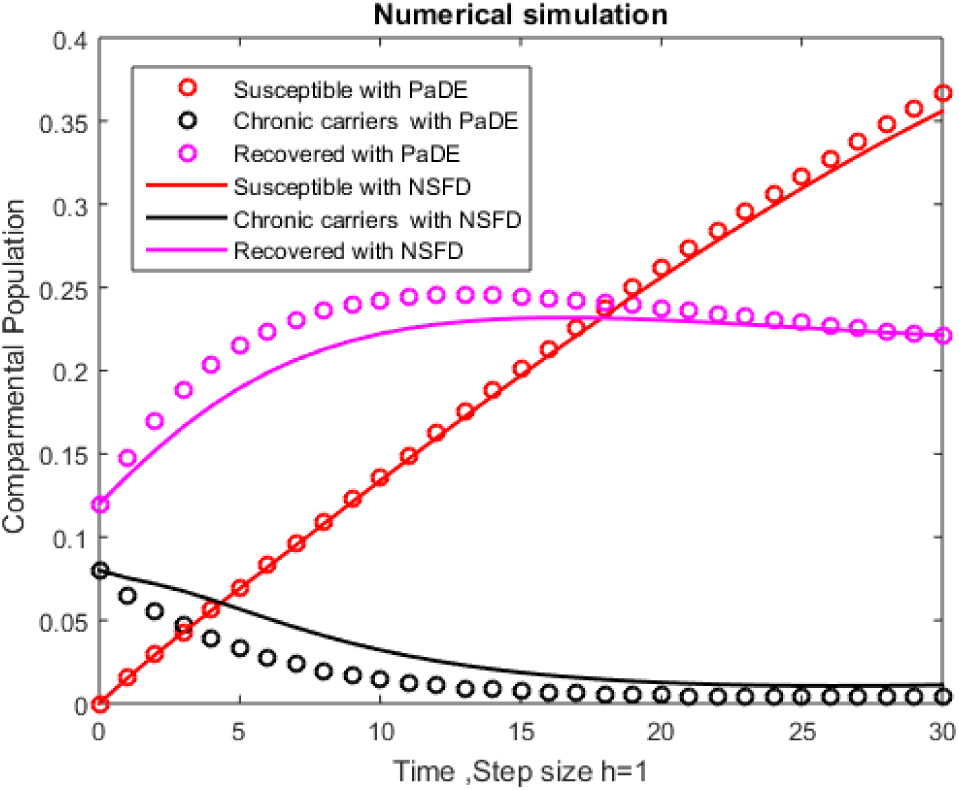
Dynamical behavior for S(t),C(t),R(t) populations in a time *t* with PaDE and NSFD.

**Figure 20.**
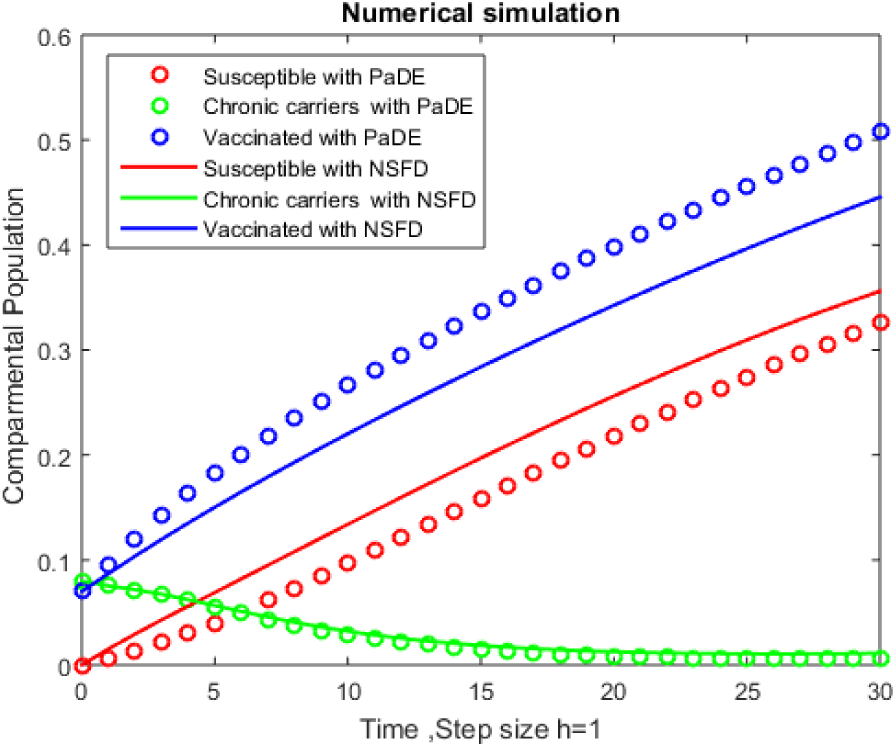
Dynamical behavior for S(t),C(t),V(t) populations in a time *t* with PaDE and NSF.

**Figure 21.**
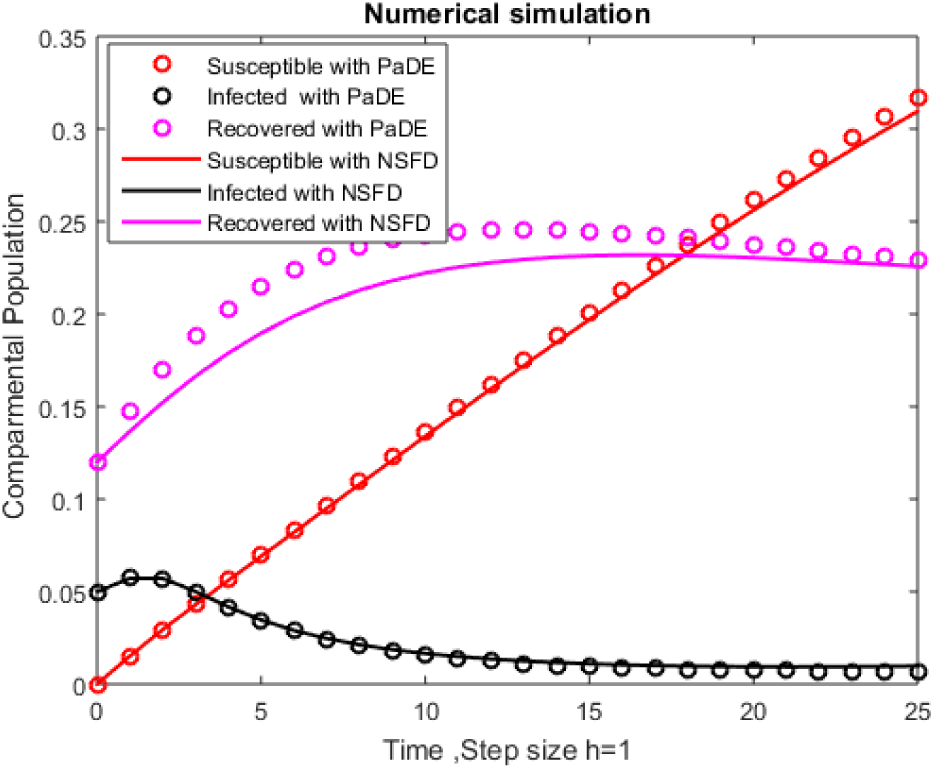
Dynamical behavior for S(t),I(t),R(t) populations in a time *t* with PaDE and NSFD.

**Figure 22.**
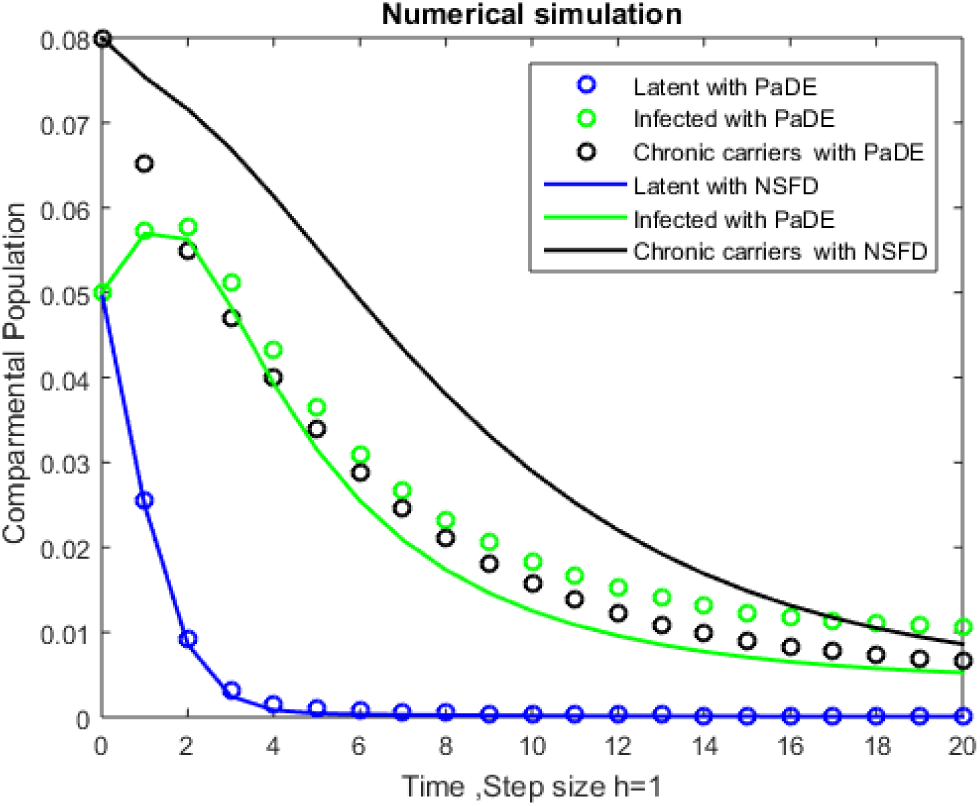
Dynamical behavior for L(t),I(t),C(t) populations in a time *t* with PaDE and NSFD.

**Figure 23.**
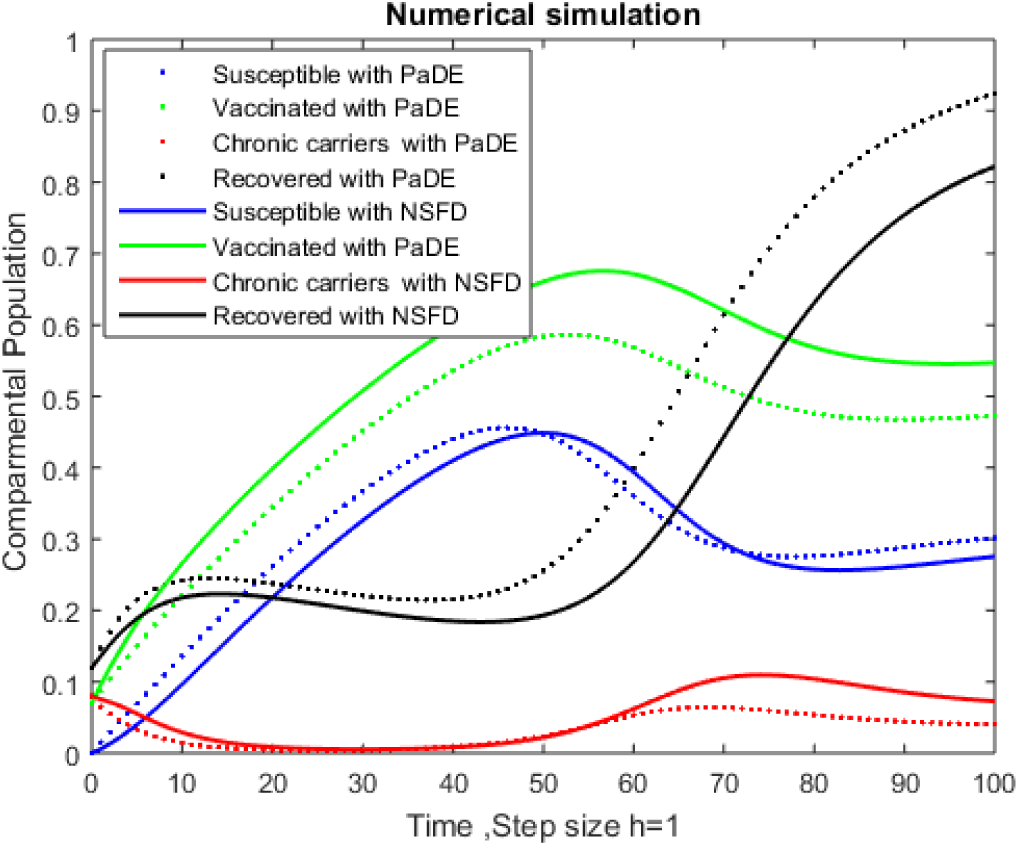
Dynamical behavior for S(t),V(t),C(t),R(t) in a time *t* with PaDE and NSFD.

**Figure 24.**
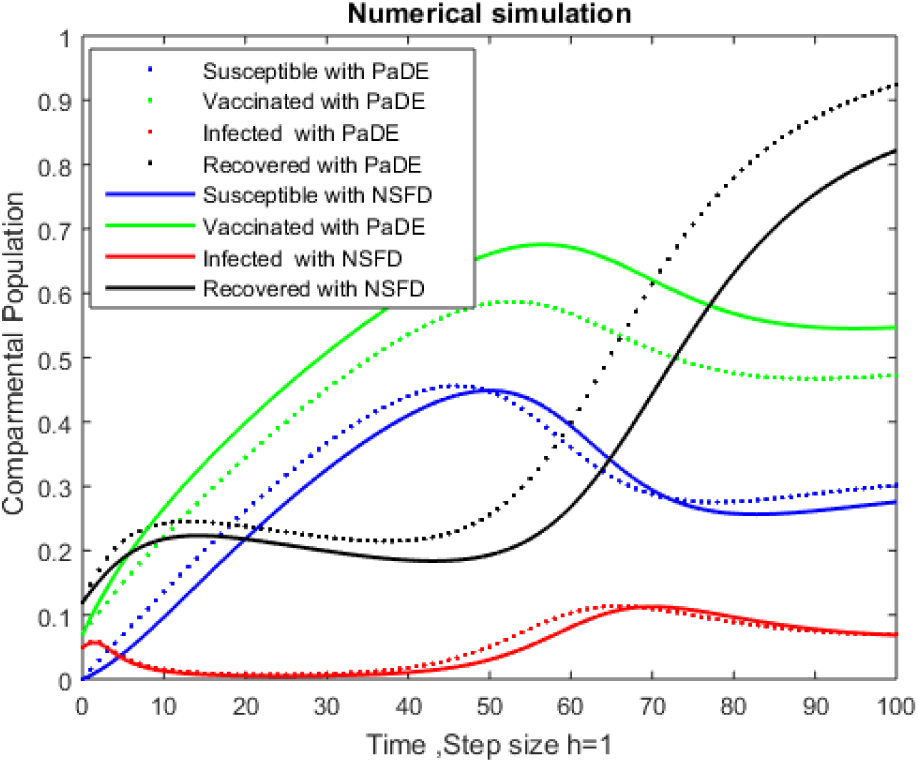
Dynamical behavior for S(t),V(t),I(t),R(t) in a time *t* with PaDE and NSFD.

**Figure 25.**
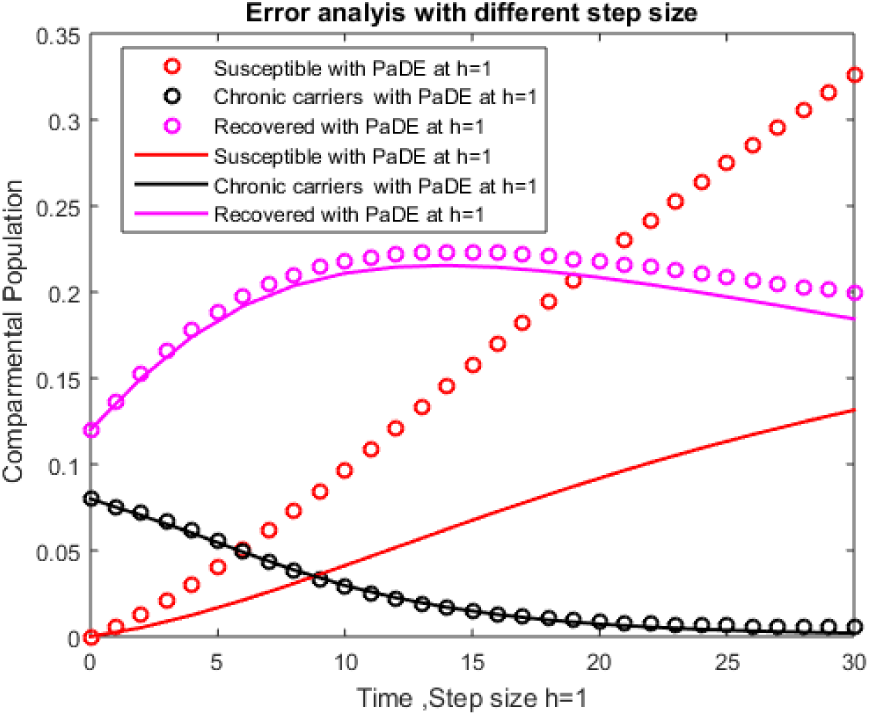
Error Analysis of S(t),C(t),R(t) to PaDE Algorithm at step size h=1,h=2.

**Figure 26.**
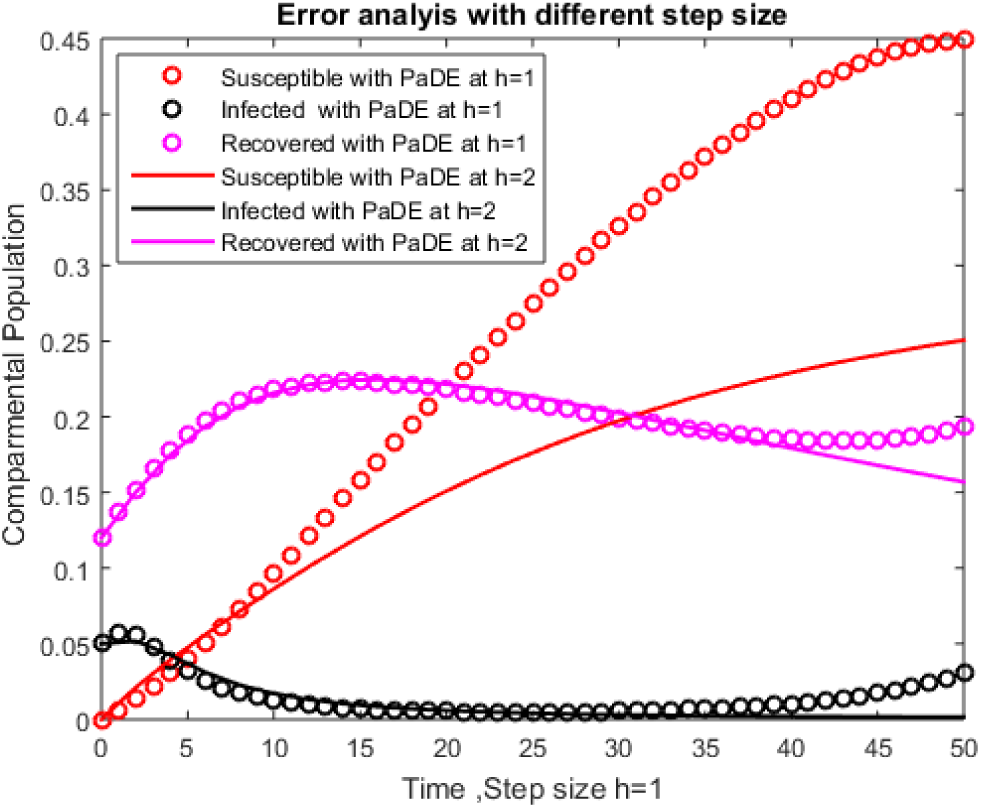
Error Analysis of S(t),I(t),R(t) to PaDE Algorithm at step size h=1,h=2.

**Figure 27.**
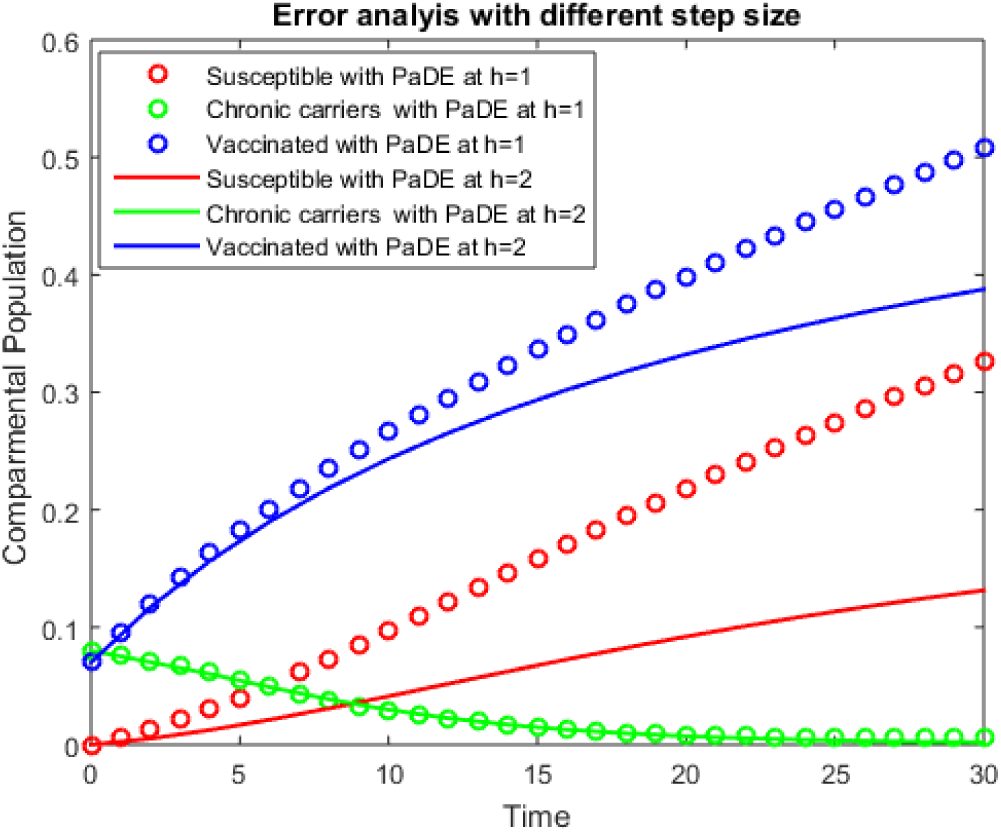
Error Analysis of S(t),C(t),V(t) to PaDE Algorithm at step size h=1,h=2.

**Figure 28.**
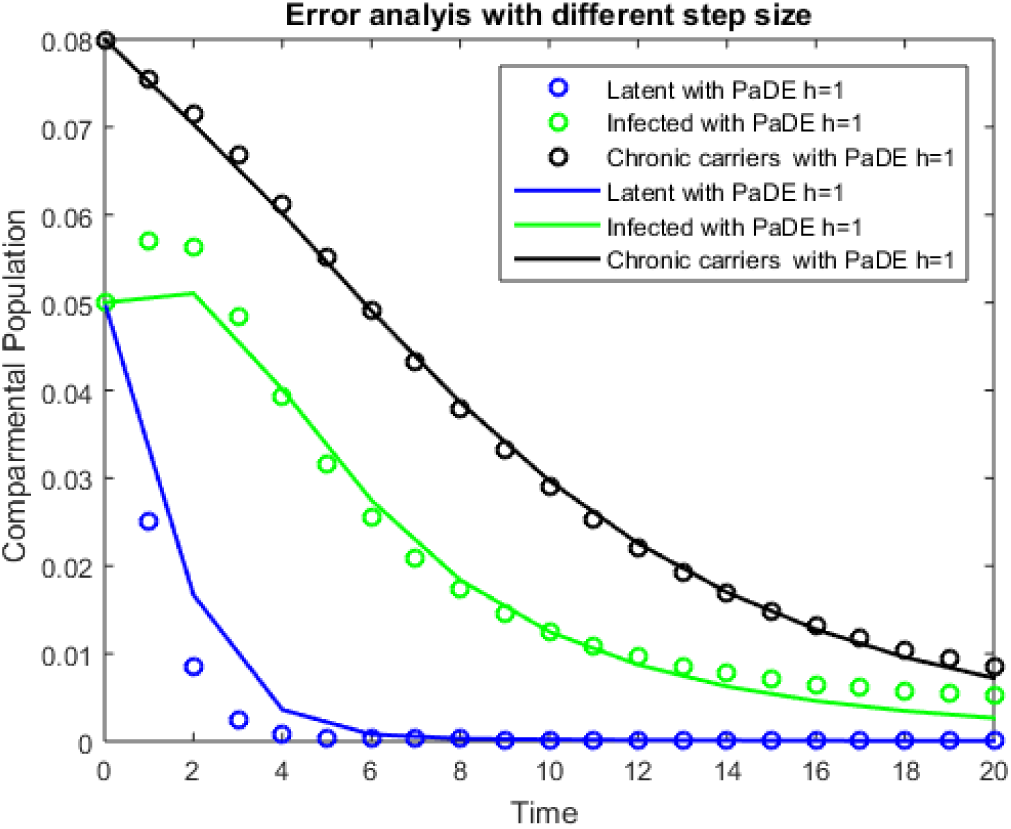
Error Analysis of L(t),I(t),C(t) to PaDE Algorithm at step size h=1,h=2.

**Figure 29.**
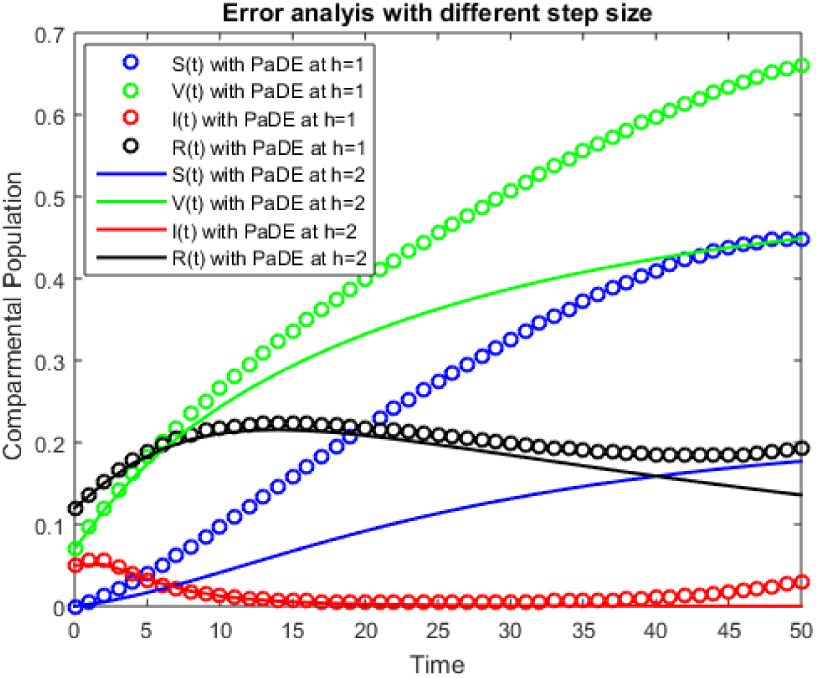
Error Analysis of S(t),V(t),I(t),R(t) to PaDE Algorithm at step size h=1,h=2.

**Figure 30.**
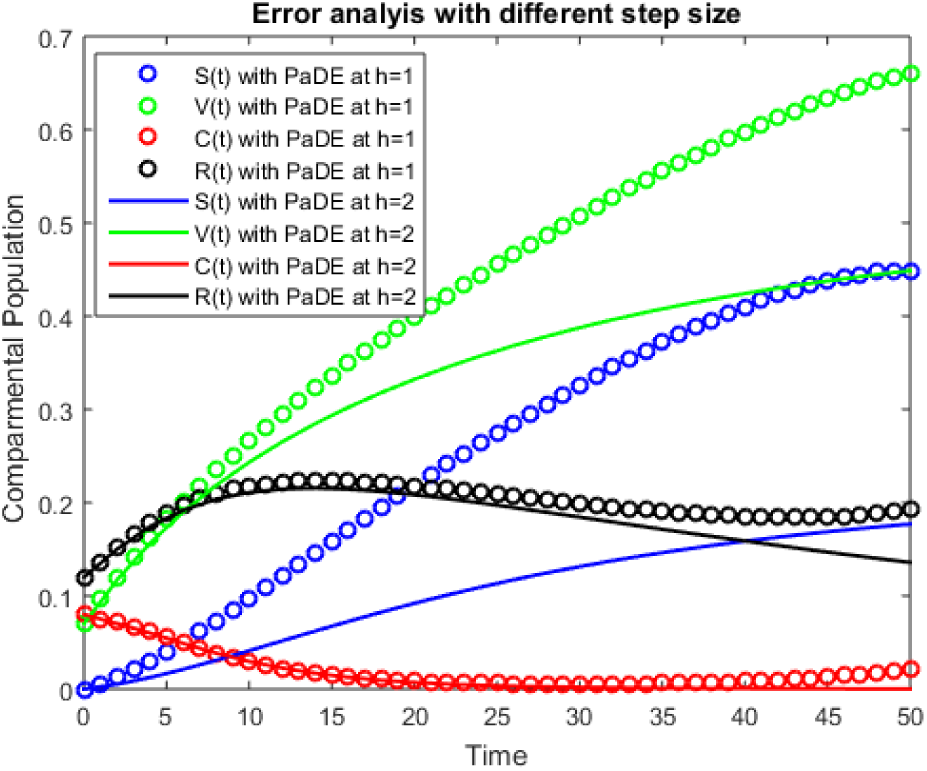
rror Analysis of S(t),V(t),C(t),R(t) to PaDE Algorithm at step size h=1,h=2.

**Table 2.** Values of physical parameters of the model.

| Parameters | Value | Parameters | Value | Parameters | Value |
| --- | --- | --- | --- | --- | --- |
| $\nu$ [41] | 0.11 | $\varphi$ [41] | 0.16 | $\mu_1$(Assumed) | 0.02 |
| $\Psi$ [41] | 0.1 | $\mu$ [45] | 0.0367 | $\gamma_3$(Assumed) | 0.1 |
| $\sigma$ [41] | 6 per year | $\mu_0$ [45] | 0.0166 | $\alpha$(Assumed) | 0.2 |
| $\beta$[46] | 0.95 | $q$ [41] | 0.885 | $\omega$(Assumed) | 0 |
| $\gamma_1$[41] | 4 per year | $\gamma_2$ [41] | 0.025 | | |

## Acknowledgements

We would like to express sincere thanks to the reviewers for their highly insightful and valuable suggestions concerning our paper.

